# The ratio of auxin to cytokinin controls leaf development and meristem initiation in *Physcomitrium patens*

**DOI:** 10.1101/2022.12.05.519125

**Authors:** Joseph Cammarata, Adrienne H. K. Roeder, Michael J. Scanlon

## Abstract

Crosstalk between auxin and cytokinin contributes to widespread developmental processes, including root and shoot meristem maintenance, phyllotaxy, and vascular patterning. Although auxin and cytokinin are potent regulators of plant development, our understanding of crosstalk between these hormones is limited to few model systems. The moss *Physcomitrium patens* (formerly *Physcomitrella patens*) is a powerful system for studying plant hormone function. Auxin and cytokinin play similar roles in regulating moss gametophore (upright shoot) architecture, as they do in flowering plant shoots. However, auxin-cytokinin crosstalk is poorly understood in moss. Here we find that the ratio of auxin to cytokinin is an important determinant of development in *P. patens*, especially during leaf development and branch stem cell initiation. Addition of high levels of auxin to *P. patens* gametophores blocks leaf outgrowth. However, simultaneous addition of high levels of both auxin and cytokinin partially restores leaf outgrowth, suggesting that the ratio of these hormones is the overriding factor. Likewise, during branch initiation and outgrowth, chemical inhibition of auxin synthesis phenocopies cytokinin application. Finally, cytokinin insensitive mutants resemble plants with altered auxin signaling and are hypersensitive to auxin. In summary, our results suggest that the ratio between auxin and cytokinin signaling is the basis for developmental decisions in the moss gametophore.

## Introduction

The hormones auxin and cytokinin are potent regulators of plant development. Their ability to specify root and shoot identity has long been exploited to regenerate plants in tissue culture^1^. In recent decades, specific roles for auxin and cytokinin have been elucidated for a wide range of developmental processes^2,3^. Each hormone elicits manifold responses depending upon cell type and developmental stage. For example, auxin can induce cell elongation and differentiation in shoots, and supports the maintenance of an undifferentiated state in the root apical meristem, whereas cytokinin promotes cell division and stem cell maintenance in shoots, but cell differentiation in roots^4–7^. Original studies of plant regeneration in culture suggested that the ratio of auxin to cytokinin was the defining factor in determining whether a given tissue would generate callus, roots, or shoots^1^. Thus, auxin and cytokinin responses depend on crosstalk between these signaling pathways.

Crosstalk between auxin and cytokinin is widespread, and is critical for varied developmental processes such as root meristem maintenance^8^, root branching architecture^9^, and shoot stem cell maintenance^6^. Crosstalk comprises many mechanisms; one hormone can modulate the transport^10–12^, synthesis^13,14^, and/or signaling pathway of the other^8^. Several of these mechanisms can occur simultaneously. For example, during root vascular patterning, auxin induces both cytokinin biosynthesis and a cytokinin signaling inhibitor, thereby increasing cytokinin response in neighboring cells (via mobile cytokinin) but not in cells with high auxin response (due to the cell autonomous cytokinin signaling inhibitor)^14^. Overall, our understanding of how auxin and cytokinin interact to regulate development is limited to the model angiosperm model plant *Arabidopsis thaliana*.

The moss *Physcomitrium patens* is a powerful model system for studying plant evolution and development^15^. Although the haploid leafy shoot (gametophore) of moss and the diploid (i.e. sporophytic) angiosperm shoot evolved independently, mounting evidence indicates that the development of both structures is accomplished by similar gene functions^16–18^. For example, *P. patens* and flowering plants exhibit auxin-mediated apical dominance, where auxin derived from the shoot tip inhibits branching at nearby nodes^19,20^.

As in Arabidopsis, auxin and cytokinin are important regulators of cell fate and of developmental transitions in *P. patens*^21^. *P. patens* begins development as a haploid spore, which gives rise to a branched network of photosynthetic filaments called protonemata. A small proportion of protonemal filaments form buds that establish a tetrahedral apical stem cell. The tetrahedral apical cell functions as the shoot apical meristem by dividing asymmetrically to yield a progenitor cell of the leaf-like phyllid and to regenerate the stem cell. Along the maturing gametophore, axillary branches are formed by the specification of new stem cells from epidermal cells in the axils of phyllids, in a manner reminiscent of axillary branching in vascular plants. In the best-understood instance of auxin-cytokinin crosstalk from moss, auxin and cytokinin synergistically promote bud formation. Auxin induces the expression of the *AINTEGUMENTA, PLETHORA* and *BABY BOOM* (*APB*) homologs, whereas cytokinin and APBs together promote the formation of buds through unknown downstream mechanisms^22^. Exogenous auxin and/or over expression of *APB* genes causes a moderate increase in bud production. Although cytokinin has a stronger effect on budding than auxin treatment or *APB* overexpression, *apb* quadruple mutants make no buds in response to cytokinin^22^, highlighting the interplay of these two hormones.

Once the gametophore is formed, auxin and cytokinin act antagonistically; these functions are analogous to those in angiosperms shoots. With regard to gametophore branch formation, auxin inhibits branch initiation, while cytokinin promotes branching^19^. Each hormone also substantially impacts phyllid development: cytokinin increases phyllid width via promoting cell division, whereas auxin promotes cell elongation and inhibits cell division, leading to the formation of long, narrow phyllids^23^. Despite the importance of auxin and cytokinin in moss gametophores, the mechanisms by which they together control branching and phyllid development are poorly understood.

Here we examine the contributions of auxin, cytokinin, and their interaction to moss gametophore development. We demonstrate that auxin and cytokinin antagonistically regulate phyllid cell differentiation and branch outgrowth. For example, reducing auxin synthesis phenocopies increased cytokinin signaling in both phyllid development and stem-cell initiation in branches. Likewise, cytokinin receptor mutants that are unable to perceive cytokinin have increased auxin sensitivity. Overall, our work demonstrates the importance of the ratio of auxin to cytokinin signaling during moss shoot development and provides a foundation for exploring auxin and cytokinin function and crosstalk outside of angiosperms.

## Results

### Cytokinin and auxin ratios regulate phyllid outgrowth and development

To test how auxin and cytokinin interact to regulate gametophore morphogenesis, we grew wild type *P. patens* on a range of concentrations of auxin and cytokinin and observed the morphology of three-week-old gametophores. Specifically, *P. patens* was grown on a three-by-three matrix of concentrations of 0 nM, 100 nM, and 2500 nM exogenous auxin (NAA), and by 0 nM, 25 nM, and 250 nM cytokinin (BAP), for a total of nine conditions to assess the impacts on gametophore development of low and high auxin and cytokinin individually and in combination (Figure 1).

**Figure 1:**
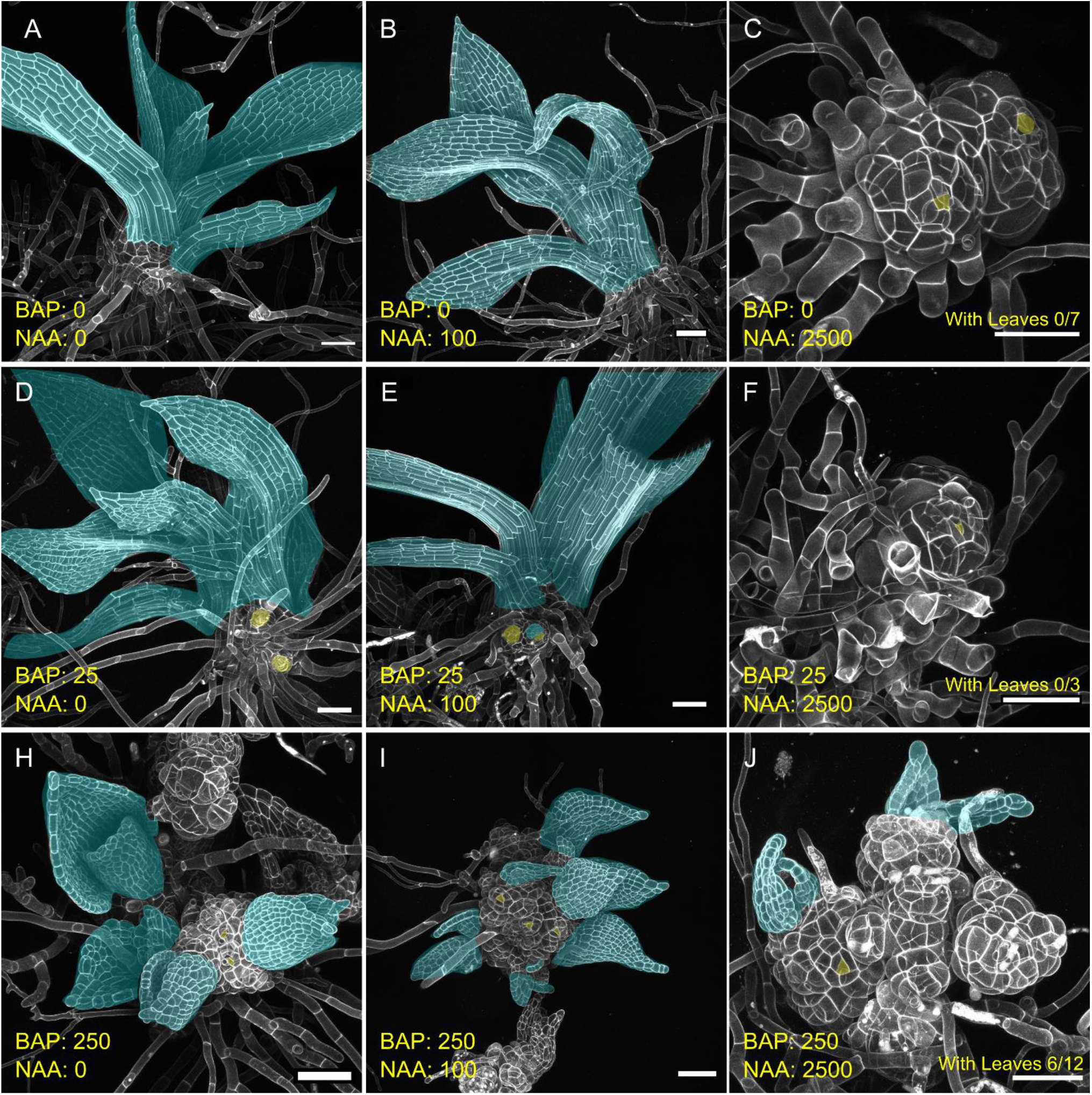
Effects of the auxin to cytokinin ratio on moss gametophore morphology. Wild type moss grown on a range of concentrations of the auxin NAA and cytokinin BAP. From left to right: 0 (A,D,H), 100 (B,E,I), and 2500 nM NAA (C, F, J). From top to bottom: 0 (A,B,C), 25 (D,E,F), and 250 nM BAP (H,I,J). Phyllids are highlighted blue. Visible or ectopic stem cells or nascent ectopic branches are highlighted yellow. Scalebars: 100 nm

Low doses of auxin (100 nM NAA) mildly impacted gametophore development, causing the formation of slightly narrow, elongated stems and phyllids (Figure 1 B). Meanwhile, increased auxin concentration (2500 nM NAA) had severe effects on development, inhibiting phyllid outgrowth and gametophore elongation while promoting rhizoid formation (Figure 1 C) as described^21,24^. In contrast, growth on low doses of cytokinin (25 nM BAP) generated wider phyllids (Figure 1 D, Supplemental Figure 1) and ectopic branch formation on stems (Figure 1 D, E). Intriguingly, gametophores grown on 25 nM BAP produced ectopic buds on rhizoids (anchoring filaments derived from gametophores), whereas cytokinin has previously been reported to induce gametophore formation on protonema (Supplemental Figure 1). Gametophores grown on high doses of cytokinin (250 nM BAP) produced wide phyllids comprising an abundance of small cells and ectopic stem cells that produced actively growing branches, consistent with a previously-described role for cytokinin in promoting cell proliferation and apical meristem identity^23,25^. Combined applications of exogenous auxin and cytokinin indicated that the ratio of these two hormones is a critical determinant of gametophore development in moss. Whereas high doses of auxin alone (2500 nM NAA) prevented phyllid outgrowth, adding high levels of both cytokinin and auxin (250 nM BAP + 2500 nM NAA) to the media restored the outgrowth of some phyllids and increased cell proliferation (Figure 1 J, Supplemental Figure 2). Phyllid outgrowth was rescued on half of the imaged gametophores (6/12). Most of these rescued phyllids appeared small and triangular, resembling juvenile phyllids^26^. We observed one phyllid comprising two files of elongated cells, representing a case where auxin-mediated repression was barely escaped (Supplemental Figure 2). Unlike higher concentrations of cytokinin (250 nM BAP), a lower concentration of cytokinin (25 nM BAP) was insufficient to rescue the phyllid outgrowth, stem elongation, or rhizoid overproduction phenotypes caused by high auxin (2500 nM NAA; Figure 1F). Gametophores on high auxin plus low cytokinin (2500 nM NAA + 25 nM cytokinin) phenocopied gametophores grown on high auxin alone (2500 nM NAA). Together these data suggest that the ratio of cytokinin to auxin is determines whether phyllids grow out or not.

Growth on 250 nM BAP induced the production of ectopic stem cells, many of which initiate phyllids, resulting in a disorganized shoot with many axes of growth (Figure 1 H). Growth on low auxin (100 nM NAA) had little to no impact stem cell production in plants cultivated on high cytokinin (250 nM BAP; Figure 1 I). Specifically, low auxin (100 nM NAA) was insufficient to inhibit branch outgrowth stimulated by either 25 nM or 250 nM BAP (Figure 1 E, I). Although fewer stem cells appear to initiate on shoots grown in high auxin and high cytokinin, the growth of these shoots is greatly inhibited (Figure 1 J) and so reduction in the absolute number of stem cells is at least partially indirect. However, more direct antagonism between auxin and cytokinin in the control of axillary meristem initiation is supported by auxin and cytokinin biosynthesis mutants generating increased and decreased branches, respectively^19^.

Overall, these data demonstrate that phenotypes caused by a given concentration of auxin can be rescued by an appropriate concentration of cytokinin, suggesting the two pathways antagonistically regulate gametophore development and impact similar developmental processes. Specifically, our data suggest that phyllid emergence and cell proliferation are responsive to the ratio of auxin to cytokinin perceived by the shoot.

### Reducing auxin synthesis phenocopies cytokinin treatment

If the ratio of auxin to cytokinin, rather than absolute concentrations, determines moss gametophore development, we reasoned that reduced auxin concentrations should yield plants resembling those treated with cytokinin. To assess the impact of reduced auxin abundance, we grew wild type moss on media supplemented with the auxin synthesis inhibitor L-Kynurinine (L-Kyn). Wild type moss grown on 10 μM L-Kyn produced dense, bushy colonies with smaller gametophores than plants grown on minimal media, similar to moss grown on low concentrations of cytokinin (BAP) (Figure 2 A, B, C). Young L-Kyn-grown gametophores had broad phyllids with reduced cell elongation, reminiscent of cytokinin-treated gametophores (Figure 2D, Figure 1). At later stages of development, L-Kyn-grown phyllids comprised elongated cells, suggesting that older plants may overcome L-Kyn-mediated auxin inhibition (Figure 2 D).

**Figure 2:**
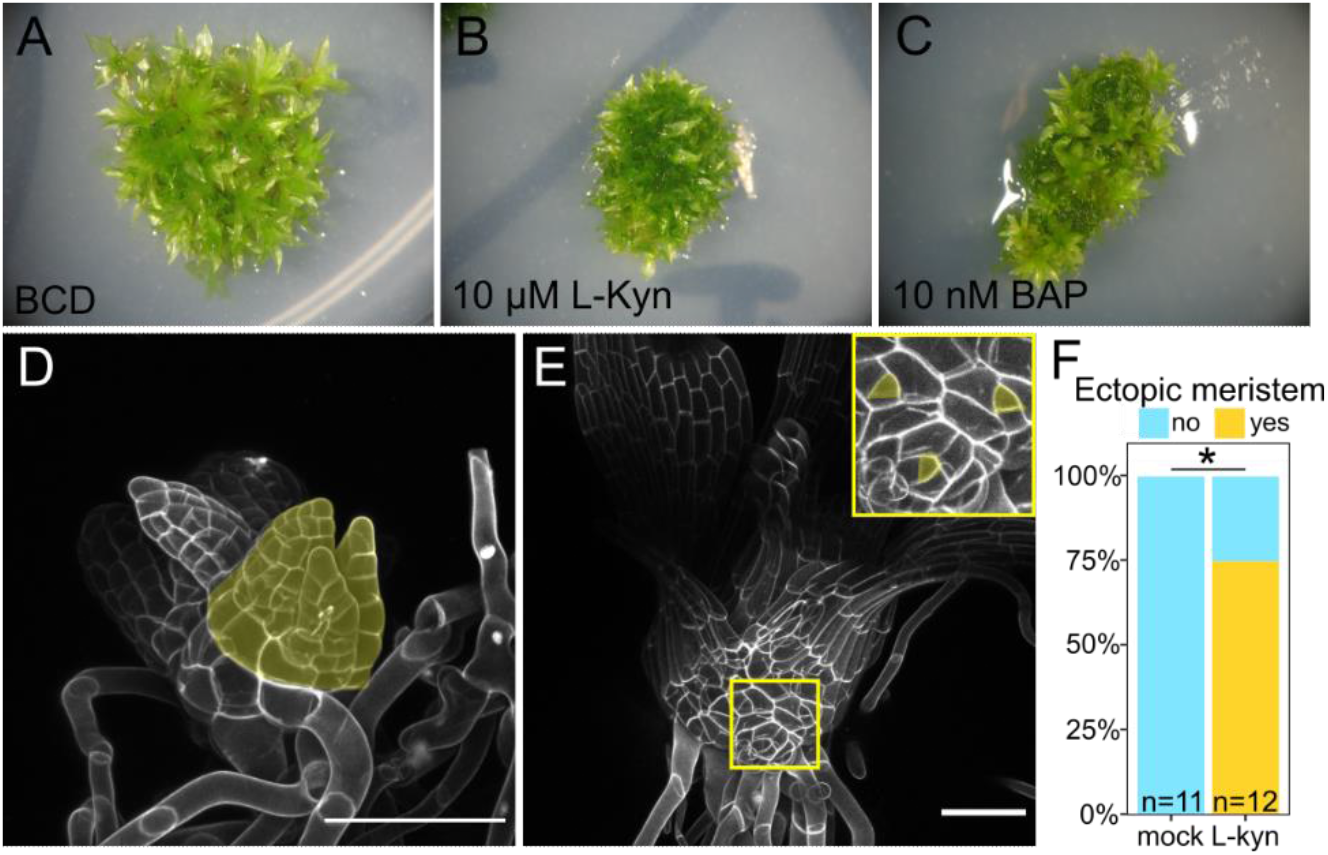
Inhibition of auxin synthesis phenocopies cytokinin treatment. Three-week-old wild type moss colonies grown in minimal media (A), on 10 μM L-Kyn (B), or on 10 nM BAP (C). A young gametophore from a moss colony grown in the presence of L-Kyn, with phyllids from an ectopic branch highlighted in yellow (D). Compare with Figure 1H. An older gametophore from moss grown with L-Kyn, with inset highlighting the presence of ectopic apical stem cells (highlighted yellow) (E). Compare with Figure 1D. Quantification of presence vs. absence of ectopic stem cell formation (F); P = 0.01, Fisher’s exact test. Scalebars: 100 nm.

L-Kyn also induced stem swelling, (Figure 2 D, E), and we detected ectopic stem cells on 75% of observed gametophores grown on 10uM L-Kyn (Figure 2F). Reduced auxin abundance correlates with increased branch number on mature gametophores^19,20^, suggesting that ectopic stem cells induced by L-Kyn are like those that give rise to axillary branches. Thus, plants with reduced auxin abundance phenocopy gametophores grown in high cytokinin concentration, consistent with the model that the ratio of auxin to cytokinin, rather than their individual absolute concentrations, determines the developmental outcome.

### Loss of cytokinin perception partially phenocopies increased auxin signalling

If our hypothesis is correct that the ratio of auxin to cytokinin concentration regulates development, we predict that loss of cytokinin perception should likewise phenocopy increased auxin. Loss-of-function triple mutants of the three cytokinin receptor-encoding *CYTOKININ HISTIDINE KINASE* genes (*CHK1, CHK2, CHK3*) are completely insensitive to exogenous cytokinin and produce small gametophores after a two week-long delay, consistent with a role for cytokinin signaling in promoting cell division and bud initiation^19,27^. Buds from *chk1 chk2 chk3* triple mutant colonies had normal division planes but smaller cells, as compared to wild type buds (Figure 3 A, D). Consistent with our hypothesis, *chk1 chk2 chk3* gametophores produced narrow phyllids with long cells and few cell files (Figure 3 E, F), a phenotype previously associated with increased auxin signaling^23,24^ (Figure 1B). In contrast, the *chk1 chk2 chk3* stems resembled plants with reduced auxin signaling. As we reported previously^25^, *chk1 chk2 chk3* gametophores exhibited increased stem cell production, contrary to expectations based on cytokinin’s role in promoting branching. Auxin represses branch outgrowth, and the increased branch formation in *chk1 chk2 chk3* triple mutants phenocopies plants with increased auxin degradation^19^. Thus, the *chk1 chk2 chk3* phenotypes suggest increased auxin signaling in the mutant phyllids (longer organs with fewer cell files) and decreased auxin signaling in the gametophore body (indicated by ectopic stem cell production). Surprisingly, *chk1 chk2 chk3* mutants exhibited a novel moss phenotype: whereas wild-type rhizoids rarely initiate branches (Figure 3 G), *chk1 chk2 chk3* mutant rhizoids initiated numerous branches that form buds, almost all of which differentiated into gametophores (Figure 3 H). Although ectopic bud formation on rhizoids has not been described in any plant where auxin synthesis, transport, or signaling are perturbed, auxin does promote bud formation^21,22^, such that ectopic bud formation in *chk1 chk2 chk3* mutants may also arise from a change in the ratio of auxin and cytokinin signaling. Overall, while the phyllid phenotypes and the formation of ectopic buds on rhizoids in *chk1 chk2 chk3* mutants support the importance of auxin-cytokinin ratios during gametophore development, ectopic stem cell formation in *chk1 chk2 chk3* triple mutants suggests a more complex regulation of stem cell initiation.

**Figure 3:**
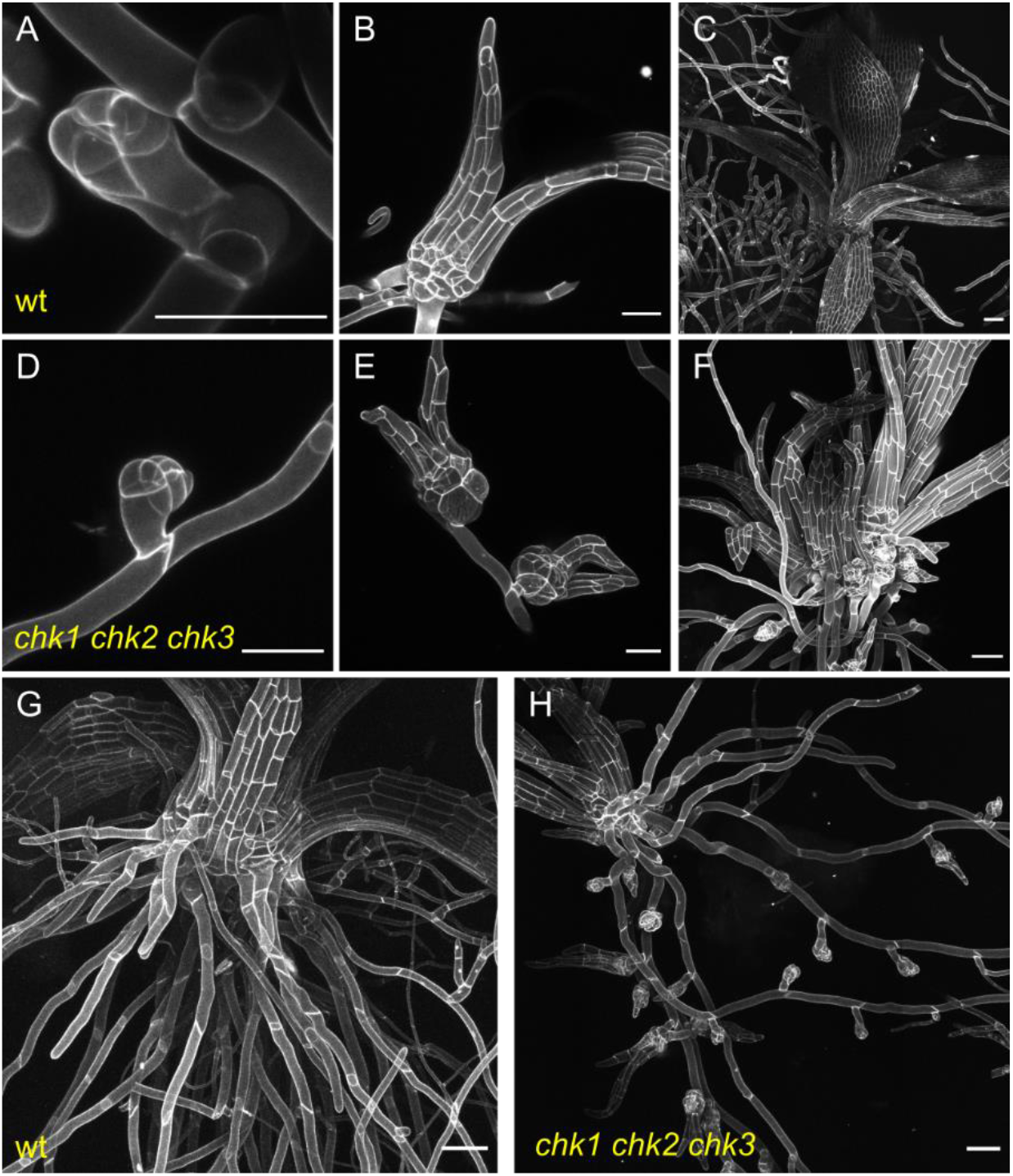
Loss of cytokinin signaling resembles altered auxin signaling, suggesting the ratio of hormones is important, but that crosstalk is complex. Wild type shoot initial (A), one-week old shoot (B), and approximately four week old shoot (C). *chk* mutant bud (D). *chk* mutant shoots have narrow leaves (E, F). *chk* shoots make ectopic branches, making them very bushy at later stages (F). Approximately four-week-old wild type (G) and *chk* (H) shoots with rhizoids, showing ectopic bud formation on *chk* rhizoids. Scalebars in A, B, D, E are 50 nm. Scalebars in C, F, G, and H are 100 nm.

### *chk* mutants are hypersensitive to auxin

Our results thus far demonstrate that reducing auxin signaling mimicked the phenotypic effects of increasing cytokinin, while reducing cytokinin mimicked the effect of increasing auxin in some organs. If it is the ratio of cytokinin to auxin that is important, we would predict that mutants with reduced cytokinin signaling will be hypersensitive to auxin application because they have no cytokinin buffer, ie. the scales are already tipped toward auxin dominating the ratio. To test whether the loss of cytokinin signaling in *chk* mutants affects responsiveness to auxin, we grew wild type and *chk1 chk2 chk3* mutants on mock media, moderate (500 nM NAA) and high (5000 nM NAA) concentrations of auxin. Wild type gametophores on 500 nM NAA had slightly enlarged stems with rounded cells (Figure 4 A). Later-staged gametophores produced a greater number of rhizoids, elongated cells in stems and phyllids, and narrow leaves (Figure 4 C). While 500 nM NAA was insufficient to suppress phyllid outgrowth in wild type gametophores, 500 nM NAA robustly suppressed phyllid outgrowth in *chk1 chk2 chk3* mutants and resulted in the formation of small gametophores with bare apical meristems and supernumerary rhizoids (Figure 4 E). These auxin-treated *chk1 chk2 chk3* gametophores resembled wild type gametophores grown on much higher concentrations of auxin (5000 nM NAA; Figure 4 B, D). These results suggest that the *chk1 chk2 chk3* triple mutant gametophores are hypersensitive to auxin, consistent with our prediction. Moreover, increasing the concentration of NAA to 5000 nM only slightly increased phenotypic severity, as compared to *chk1 chk2 chk3* grown on 500 nM NAA. Specifically, we sometimes observed small cellular projections suggesting that phyllid initial identity was attained, but subsequent development was rapidly terminated on 500 nM NAA. No small phyllid initials were identified following treatment with 5000 nM NAA (Supplemental Figure 3, Figure 4 F). Therefore, not only are *chk1 chk2 chk3* mutants hypersensitive to auxin but also, the auxin response appears to saturate. These findings are consistent with the ratiometric model of auxin-cytokinin interaction: in *chk1 chk2 chk3* bereft of cytokinin signaling, a lower concentration of auxin is required to tip the scales to inhibit phyllid outgrowth than in wild type gametophores.

**Figure 4:**
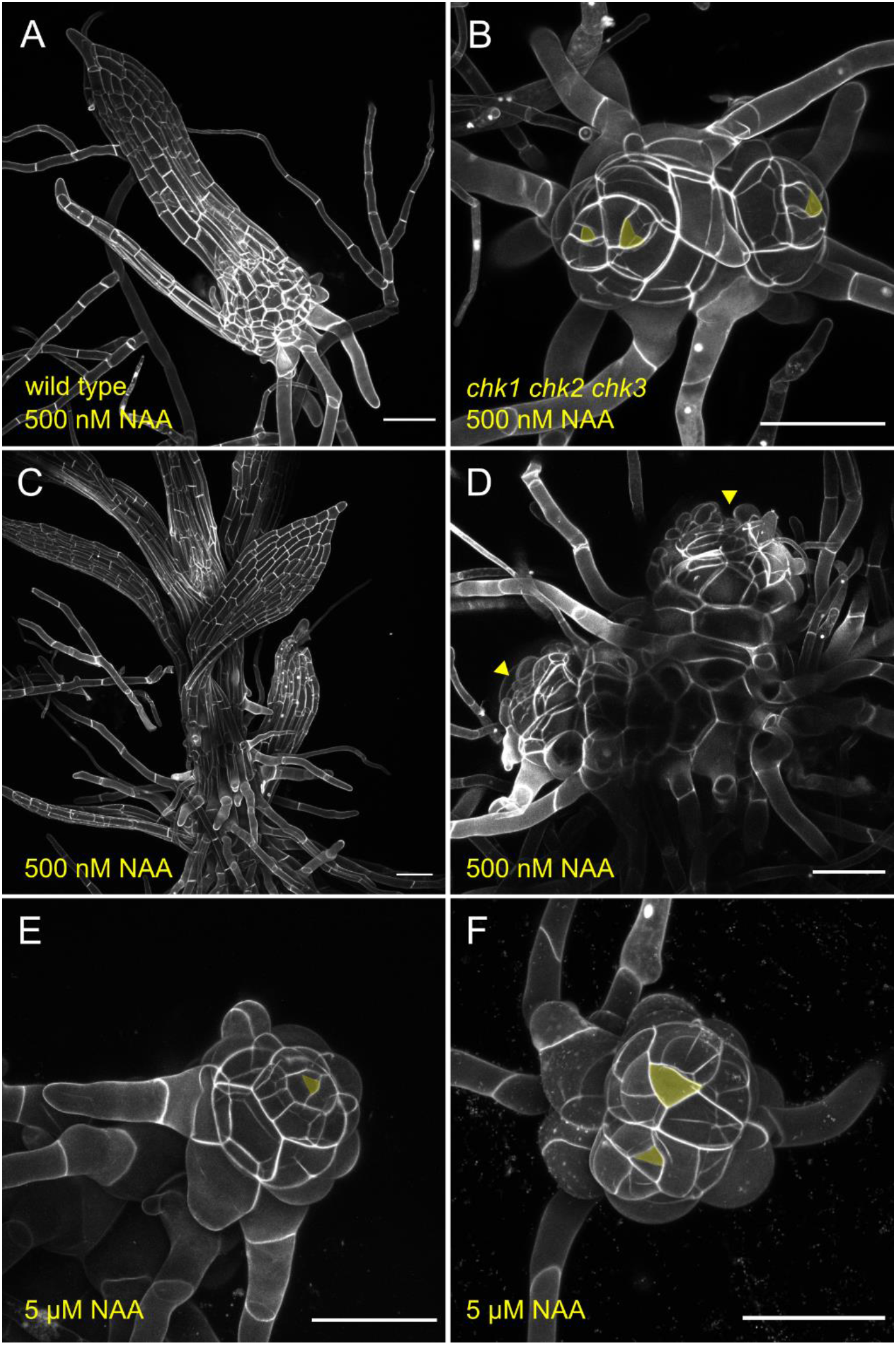
*chk* mutants are hypersensitive to auxin. Early (A, B) and late (C, D) stage wild type (A, C) and *chk* (B, D) plants grown on 500 nM NAA. Whereas 500 nM NAA causes cell expansion and the formation of elongated stems and leaves in wild type moss, *chk1 chk2 chk3* mutants on 500 nM NAA resemble wild type moss grown on 5000 nM NAA. Wild type (E) and *chk1 chk2 chk3* (F) shoots grown on 5000 nM NAA have bare apices with no leaf emergence and increased rhizoid production. Scalebars: 100 nm.

Whereas the ratio of auxin to cytokinin signaling was critical for phyllid development, the regulation of branching by auxin and cytokinin appeared to be more complex. We often observed multiple stem cells on *chk1 chk2 chk3* mutants grown on auxin, suggesting that 500nM NAA is sufficient to inhibit phyllid outgrowth in these triple mutants but not the formation of ectopic stem cells. However, *chk1 chk2 chk3* mutant gametophores still formed multiple stem cells in the presence of 5000 nM NAA (Figure 5 F). Thus, even a very high auxin concentration was insufficient to prevent ectopic SAM formation in *chk1 chk2 chk3*. These data suggest that the mechanism(s) whereby auxin inhibits stem cell formation is impaired in *chk1 chk2 chk3* mutants and that, unlike during phyllid development, the control of branch development did not hinge on the ratio of auxin to cytokinin signaling.

## Discussion

We examined the effects of auxin and cytokinin concentration, alone and in combination, on moss gametophore development. In general, the ratio of auxin to cytokinin regulates phyllid cell proliferation, and branch formation, with auxin and cytokinin acting antagonistically. Reducing auxin synthesis phenocopies cytokinin treatment, while cytokinin receptor mutants partly phenocopy auxin treatment. Thus, the ratio of auxin to cytokinin signaling is critical for directing gametophore development. However, high concentrations of auxin were not sufficient to inhibit ectopic meristem initiation in *chk1 chk2 chk3* mutants, which are defective in cytokinin signaling. These data suggest that auxin modulates stem cell formation in a cytokinin-dependent manner and supports our previous hypothesis that cytokinin is upstream of a separate pathway regulating SAM initiation^25^. Our work supports the longstanding concept that the ratio of auxin to cytokinin, not the concentration of either independently, regulates plant development, in a new and evolutionary distant lineage of plant from where this concept was generated.

### Antagonistic control of phyllid development

High concentrations of exogenous auxin can inhibit phyllid outgrowth, and the dose of auxin at which phyllid outgrowth is robustly inhibited has previously been used as a gauge of auxin sensitivity^21^. We found that phyllid outgrowth in *chk1 chk2 chk3* mutants was hypersensitive to auxin. A moderate concentration of NAA (500 nM) suppressed phyllid outgrowth in *chk1 chk2 chk3* mutant but not wild type gametophores. Inversely, supplementing plants with a high concentration of cytokinin rescued auxin-mediated inhibition of phyllid outgrowth. Therefore, auxin and cytokinin act antagonistically to regulate phyllid development, and the ratio of these two hormones (not their absolute values) is the determining factor.

We speculate that two, non-exclusive mechanisms explain the antagonistic regulation of phyllid outgrowth; both explanations are consistent with the hypothesis that the ratio of auxin to cytokinin signaling is critical for phyllid development. First, auxin and cytokinin might independently regulate the potential of the phyllid initial cell to divide. After the asymmetric division of the stem cell to produce a daughter destined to form a phyllid, the phyllid initial undergoes repeated, self-renewing cell divisions that produce cells for early phyllid development^26^. In *chk1 chk2 chk3* mutants, phyllids are composed of fewer files of longer cells, indicating a reduced capacity to divide (Figure 1)^27^. Similar phenotypes have been shown for *P. patens* treated with auxin^21,23,24^. Thus, the inhibition of phyllids on *chk1 chk2 chk3* gametophores grown on auxin could represent the cumulation of two independent mechanisms that reduce cell division competency. Likewise, rescued phyllid outgrowth observed after combining 250 nM BAP and 2500 nM NAA would result from cytokinin promoting division of the phyllid initial just long enough to generate a few files of phyllid cells before division terminates. In support of this idea, not all gametophores on 250 nM BAP and 2500 nM NAA escape phyllid suppression, and some form small leaves with as few as two cell files (Supplemental Figure 2).

A second possibility is that cytokinin might alter auxin signaling and response at the cellular level, and vice versa. A similar interaction between cytokinin and auxin regulates the size of the root meristem, where cytokinin induces the expression of *AUX/IAA* genes that encode transcriptional repressors of auxin signaling^4^. In moss filaments, cytokinin similarly promotes *AUX/IAA* expression^28^. As yet, the transcriptional response of moss gametophores to cytokinin has not been assessed. Our data demonstrating the antagonistic effects of auxin and cytokinin on phyllid outgrowth and development are consistent with a role for cytokinin in reducing the sensitivity of cells to auxin. In this scenario, *chk1 chk2 chk3* mutants would be hypersensitive to auxin, explaining the high-auxin phenotype when grown on minimal media and the full repression of phyllid outgrowth at 500 nM NAA. Exogenous BAP would then be expected to reduce sensitivity of cells to auxin, explaining the re-emergence of leaves on gametophores grown on both high cytokinin and high auxin.

### Auxin modulates stem cell formation in a cytokinin-dependent manner

Auxin and cytokinin antagonistically regulate branch formation in moss. Growth on the auxin synthesis inhibitor L-Kyn resulted in gametophores with ectopic branches and supernumerary shoot meristems even at early developmental stages, demonstrating the importance of auxin during inhibition of axillary SAM initiation and outgrowth.

Although cytokinin promotes meristem formation, *chk1 chk2 chk3* mutants that are devoid of cytokinin perception counter-intuitively form ectopic meristems. We have previously shown that a network where cytokinin promotes stem cell formation via a pathway buffered by an incoherent feedforward loop can explain stem cell specification rates in wild type, as well as in stem cell homeostasis mutants grown on a range of cytokinin concentrations^25^. However, the role of auxin in this proposed pathway is unclear. Reducing auxin synthesis with L-Kyn led to the formation of ectopic stem cells, swollen stems, and broad phyllids, phenocopying plants treated with cytokinin. These data suggest that auxin and cytokinin interact antagonistically to regulate meristem specification in moss. Because reduced auxin signaling and exogenous cytokinin elicit similar effects, again it appears that the ratio and not the absolute amount of signaling through each pathway is important.

Triple mutant *chk1 chk2 chk3* moss forms long, narrow phyllids suggesting increased auxin response, and initiate ectopic stem cells, suggesting decreased auxin response within the same plant. One explanation for this differential change in auxin response in these two tissues is an altered auxin distribution in *chk1 chk2 chk3* mutants. In *Arabidopsis thaliana*, cytokinin signaling controls the directionality and abundance of the PIN family of polar auxin efflux carriers^7,12,29,30^; this interaction may also exist in moss. However, very high concentrations of auxin were unable to suppress ectopic branch formation in *chk1 chk2 chk3* mutants, suggesting that redistributed auxin alone does not explain ectopic branch formation in *chk1 chk2 chk3*, and a different, auxin-independent mechanism is at play during ectopic stem cell production in these triple mutants. Therefore, it is still possible that auxin and cytokinin exert control over the same phenotypes through distinct pathways.

## Supplemental Figures

**Supplemental Figure 1:**
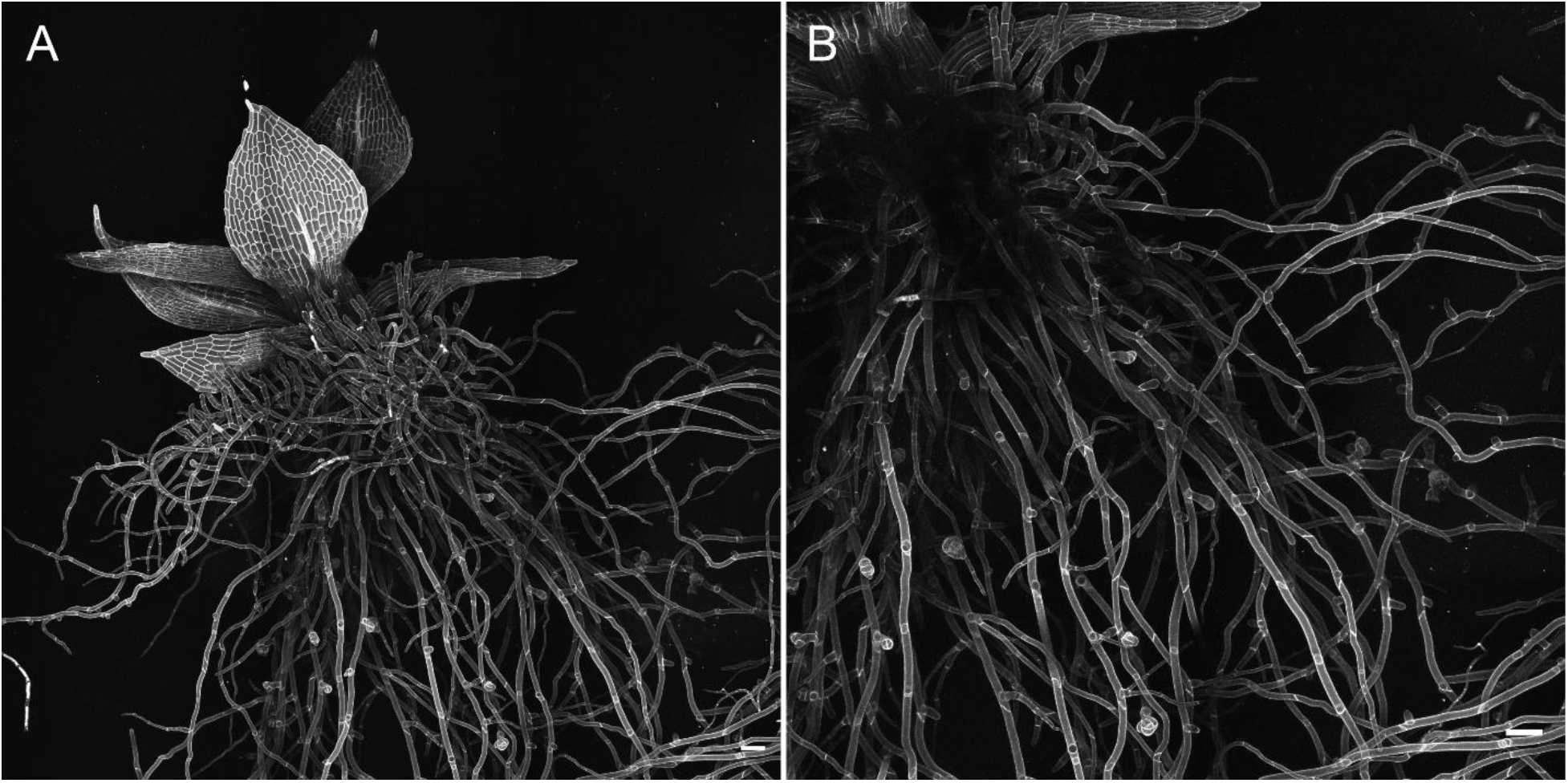
Treatment with 25 nM BAP causes leaf widening and ectopic bud formation on rhizoids. A) Z-projection of a shoot grown on 25 nM BAP. B) projection of a smaller area and fewer Z-slices of the image in panel A to better show rhizoids and ectopic buds. Scale bars: 100 nm

**Supplemental Figure 2:**
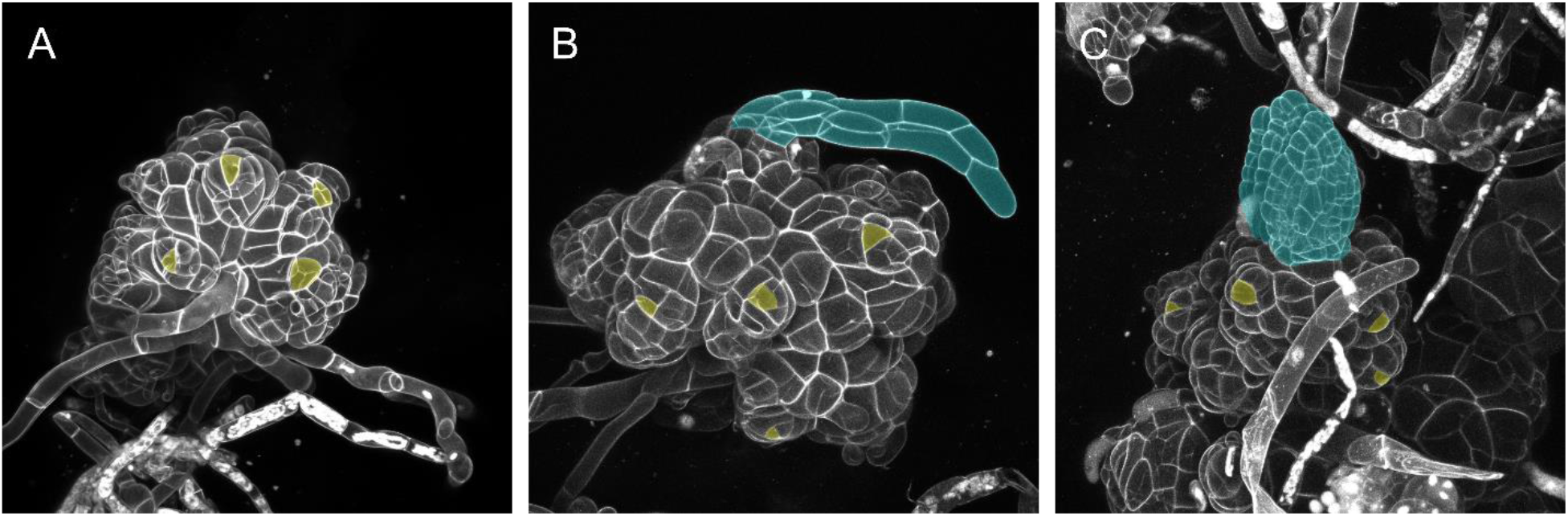
High concentrations of cytokinin variably rescue leaf-suppression caused by high concentrations of auxin. Each image depicts shoots grown on 250 nM BAP and 2500 nM NAA. Suppression of leaves can be complete (A), partial (B), or abrogated (C). Leaves pseudo colored blue and visible apical stem cells pseudo colored yellow.

**Supplemental Figure 3:**
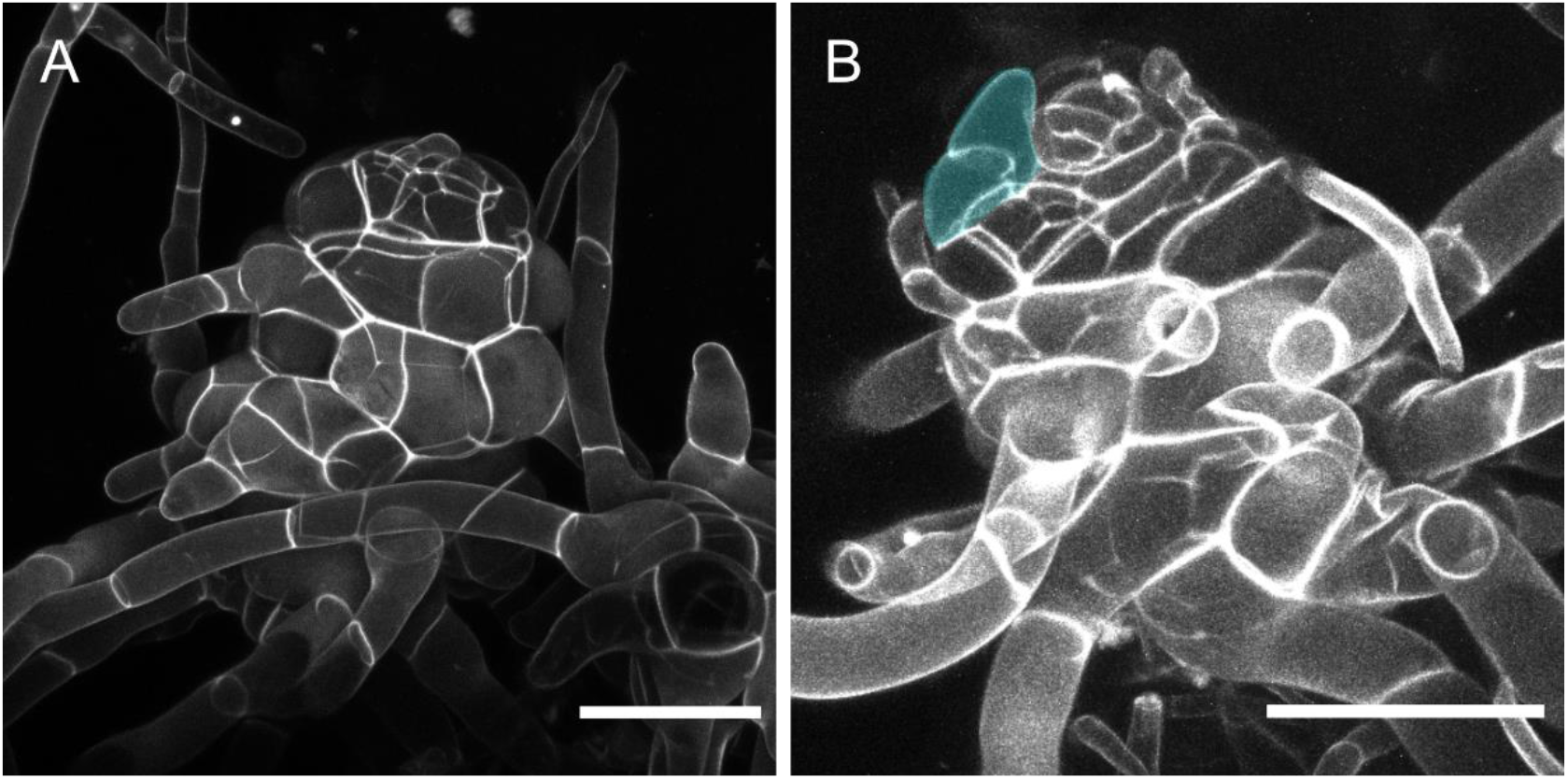
*chk* mutants are hypersensitive to exogenous NAA. Two shoots grown on 500 nM NAA, a concentration insufficient to inhibit shoot elongation or leaf emergence in wild type. Leaf formation is completely inhibited in *chk* shoots grown on 500 nM NAA (A). Sometimes, small projections resembling one or two leaf cells can be seen (B).

## Materials and methods

### Moss culture and hormone treatment

Routine moss culture was conducted on minimal media supplemented with ammonium tartrate (BCDAT, see Media section) and overlain with cellophane. To propagate moss, tissue was blended in 5-7 ml sterile Water using a Dremel with a custom-made propeller blade attachment. 1-2 ml of blended moss was pipetted to inoculate fresh plates. All experiments and lines were of the Gransden strain of *Physcomitrium patens* (previously *Physcomitrella patens*). For phenotyping, small tufts of moss filaments growing on BCDAT were placed on agar BCD plates and grown for three to five weeks. All moss was grown under continuous light at 25 degrees Celsius. For hormone and pharmacological treatments, hormone stocks were diluted to specified concentrations in BCD media and plants were grown continuously in the presence of hormone or in mock (solvent) control media.

### Confocal Microscopy and Phenotypic Analysis

Moss gametophores for imaging were dissected from three-to-five-week-old colonies grown on minimal media (BCD). Colonies were flooded with water prior to dissection and tapped vigorously to liberate air bubbles. Dissected gametophores were placed on a slide with a well made of vacuum grease and filled with 5 μg/ml propidium iodide (PI) in waer. Slides were then sealed with a coverslip and incubated for at least fifteen minutes before imaging. All confocal imaging was conducted using a Zeiss LSM-710 laser scanning microscope. PI-stained tissues were imaged using a 514 nm laser for excitation. Emission wavelengths between 566 and 650 nm were collected. Ectopic stem cells were identified on the basis of their tetrahedral shape.

#### Media

##### BCDAT

- 250mg/L MgSO_4_.7H_2_O
- 250mg/L KH_2_PO_4_ (pH6.5)
- 1010mg/L KNO_3_, 12.5mg/L
- FeSO_4_.7H_2_O
- 0.001% Trace Element Solution*
- 0.92 g/L C_4_H_12_N_2_O_6_ (ammonium tartrate)
- 8g/L agar
- CaCl_2_ added to a 1mM concentration after autoclaving.

Minimal Media (BCD) is BCDAT without the ammonium tartrate

*Trace Element Solution

- 0.614mg/L H_3_BO_3_
- 0.055mg/L AlK(SO4)2.12H_2_O
- 0.055mg/L CuSO_4_.5H_2_O
- 0.028mg/L KBr
- 0.028mg/L LiCl
- 0.389mg/L MnCl_2_.4H_2_O
- 0.055mg/L CoCl_2_.6H_2_O
- 0.055mg/L ZnSO_4_.7H_2_O
- 0.028mg/L KI
- 0.028mg/L SnCl_2_.2H_2_O

